# Retention Time Standardization and Registration (RTStaR): An algorithm that matches corresponding and identifies unique species in nanoliquid chromatography-nanoelectrospray ionization-mass spectrometry lipidomic datasets

**DOI:** 10.1101/2021.05.01.442266

**Authors:** Alexandre P. Blanchard, Yun Wang, Graeme P. Taylor, Matthew W. Granger, Stephen Fai, Daniel Figeys, Tomas Paus, SYS Consortium, Zdenka Pausova, Hongbin Xu, Steffany A.L. Bennett

## Abstract

Bioinformatic tools capable of registering, rapidly and reproducibly, large numbers of nanoliquid chromatography-nanoelectrospray ionization-tandem mass spectrometry (nLC-nESI-MS/MS) lipidomic datasets are lacking. We provide here a freely available Retention Time Standardization and Registration (RTStaR) algorithm that aligns nLC-nESI-MS/MS spectra within a single dataset and compares these aligned retention times across multiple datasets. This two-step calibration matches corresponding and identifies unique lipid species in different lipidomes from different matrices and organisms. RTStaR was developed using a population-based study of 1001 human serum samples composed of 71 distinct glycerophosphocholine metabolites comprising a total of 68,572 analytes. Platform and matrix independence were validated using different MS instruments, nLC methodologies, and mammalian lipidomes. The complete algorithm is packaged in two modular ExcelTM workbook templates for easy implementation. RTStaR is freely available from the India Taylor Lipidomics Research Platform http://www.neurolipidomics.ca/rtstar/rtstar.html. Technical support is provided through

## 1 Introduction

Lipidomics refers to the systems-level analysis of lipid diversity, lipid interactions, and lipid regulation enabled, in part, by the application of high performance liquid chromatography, electrospray ionization-tandem mass spectrometry LC-ESI-MS/MS to the profiling of lipid composition (Bennett, et al., 2013; Brown, 2012; Brown and Murphy, 2009). These profiles are revealing intriguing metabolic changes associated with aging, sex, and neurodegenerative diseases (Mapstone, et al., 2014; Rappley, et al., 2009; Ryan, et al., 2009). Comparing organismal, cellular, and subcellular organelle lipidomes, however, is difficult. One of the challenges is the limited number of bioinformatic tools capable of matching corresponding and identifying unique lipid species across lipidomes in different matrices. It remains labour-intensive for different laboratories, and even within targeted lipidomic experiments from the same laboratory, to register, rapidly and reproducibly, large numbers of MS spectra. Despite application of identical chromatographic methods, small run-to-run variations in flow rate, composition of the mobile phase, temperature, pH, packing of chromatographic columns etc., can cause substantial variations in retention time (RT) (Smith, et al., 2015). These differences are magnified when nanoLC and nanoESI-MS/MS are used. nLC-nESI-MS/MS enhances measurement sensitivity and dynamic range of low abundance, biologically potent lipid second messengers (Danne-Rasche, et al., 2018; Granger, et al., 2019; Sliz, et al., 2020; Whitehead, et al., 2007), yet is more subject to variations in RT than LC or ultraperformance LC (UPLC) making peak selection in selected reaction monitoring or multiple reaction monitor modes (SRM or MRM) across lipidomes challenging. Easily implemented algorithms capable of standardizing RTs across nLC-nESI-MS/MS datasets would facilitate this process.

Here, we describe development and validation of the <u>R</u>etention <u>T</u>ime <u>St</u>andardization <u>a</u>nd <u>R</u>egistration (RTStaR) algorithm. RTStaR was designed to (a) verify user alignment of the RTs of lipid species from run-to-run within a single dataset/lipidome, (b) standardize RT alignments across multiple runs between datasets/lipidomes, (c) predict the empirical RTs of known lipid species in any given spectra, and (d) register corresponding and unique lipids across datasets/lipidomes acquired by nLC-nESI-MS/MS. Users can also build their own nLC-nESI-MS/MS analyte databases specific to their particular nLC methodology. The complete algorithm is packaged in two modular Excel^™^ templates for easy implementation and is available online at: http://www.neurolipidomics.ca/rtstar/rtstar.html.

## 2 METHODS

### 2.1 Datasets

RTStaR was developed using three glycerophosphocholine (GPC) metabolite datasets composed of 1061 MS spectra. These datasets were: (1) a population-based study of circulating GPC metabolites in human serum consisting of 1001 participants, (2) a genotype and intervention comparison study of GPC metabolites in the hippocampus of 30 wildtype (NonTg) and N5 TgCRND8 (Tg) mice, a sexually dimorphic mouse model of Alzheimer’s disease (Granger, et al., 2016; Wang, et al., 2013), and (3) a comparative biomarker study of the plasma lipidome in these same 30 animals. To examine platform independence, RTStaR was challenged with additional datasets profiled using a different nLC methodology and MS. These validation lipidomes examined GPC metabolism in: (1) a longitudinal study from birth to elderly in the parietal-temporal cortex of 50 NonTg and Tg littermates, (2) a genetic ablation study in the parietal-temporal cortex of 49 platelet activating factor receptor (PAFR) wild-type (NonTgPAFR^+/+^), null mutant (NonTgPAFR^-/-^), TgPAFR^+/+^ and TgPAFR^-/-^ N5 TgCRND8 mice, and (3) a post-mortem study profiling the composition of GPC metabolites in the parietal-temporal cortex of 10 Alzheimer’s disease and age-matched human controls. Lipids were extracted from serum, plasma, hippocampi, and parietal-temporal cortex as we have described (Xu, et al., 2013). Accuracy was assessed using a dataset composed of 20 lipid standards (Avanti Polar Lipid and Cayman Chemical, Supplementary Data). Complete NLC-NESI-MS/MS methodologies and lipid standards details are provided in Supplementary Data.

## 3 ALGORITHM

The algorithm is based on five assumptions: (1) a specific lipid species will *theoretically* have the same RT in every nLC-nESI-MS/MS spectra if separated using the same nLC method; (2) the RT of a peak is measured at the point of maximal intensity; (3) isobaric lipids can only be distinguished if the difference in RT between peaks is measurable; and (5) empirical RT differences between runs resulting from *system-level variations* can be corrected using a monotonic function whereas *component-level variations* cannot. Following Smith et al. (2015), we define system-level variations as run-to-run differences that affect the chromatography of *all* species in a given run. These differences can result from sample extraction and preparation by multiple researchers, replacement of connection tubing between runs, fluctuations in separation column temperature, decline in column performance over time, column-to-column variations upon replacement, etc. Conversely, component-level variations are alterations that affect discontinuous chromatographic events (Smith, et al., 2015). These changes can result from transient column clogging, transient changes in flow rate, random fluctuations in ionization efficiency, periodic ion suppression, etc.

### 3.1 Registering the RT of each lipid species in nLC-nESI-MS/MS spectra

To align spectra, RTStaR begins with a user-defined calibrator dataset that (a) estimates the average system-level variation associated with the user’s particular LC methodology and (b) calculates the theoretical RT for every species present in a given lipidome (Fig. 1A). Next, it standardizes other datasets collected using identical LC methods to this calibrator dataset (Fig. 1B). Here, datasets are defined as a single study made up of a series of biological replicates, each represented by a unique MS run. A dataset can include both control and experimental groups. Our alignment and standardization pipeline is presented in Figure 1.

**Fig. 1.**
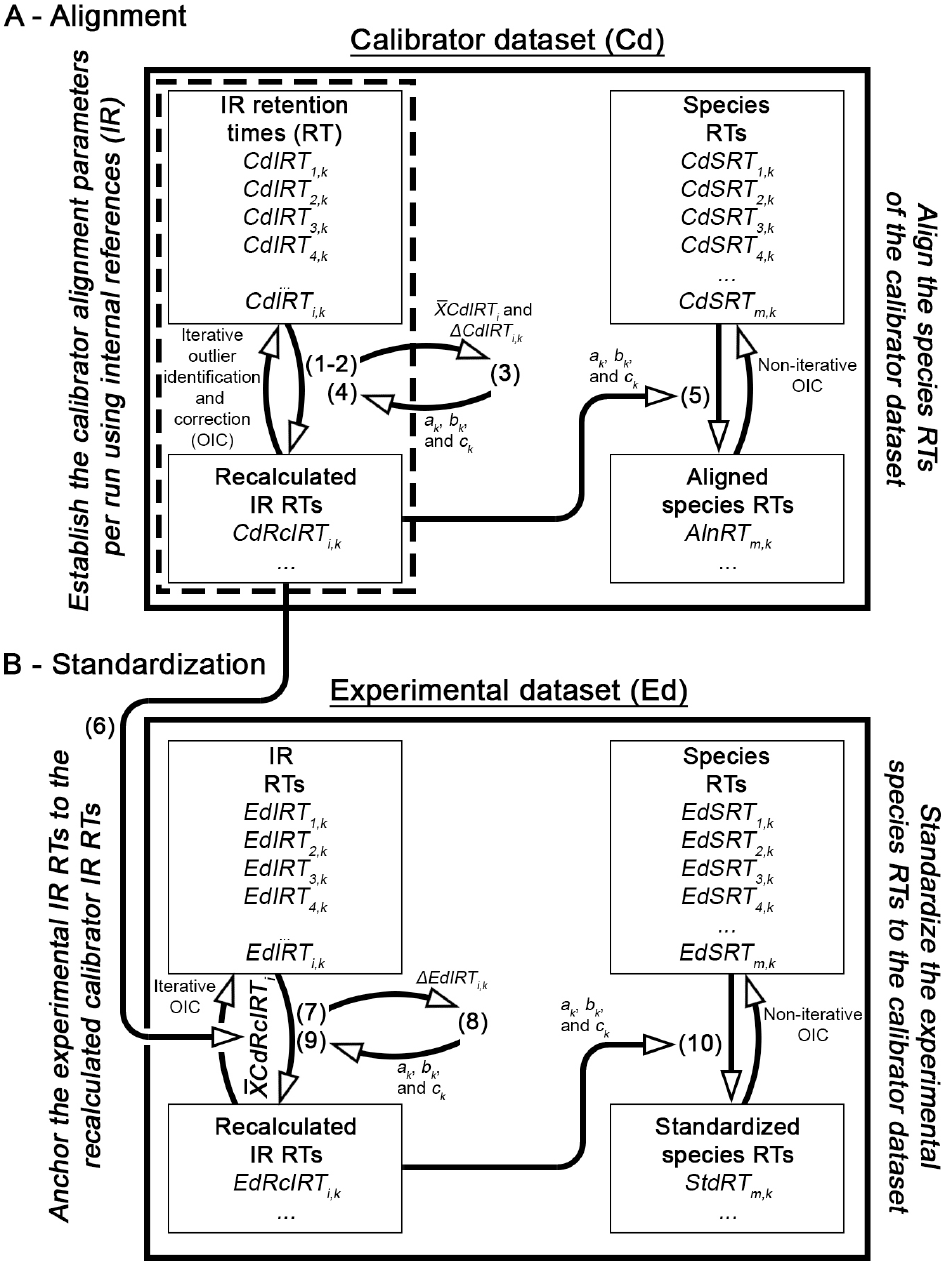
Schematic of the RTStaR spectral alignment, standardization, and registration process. Diagram depicts the iterative algorithmic pipeline through which the RTs of species in multiple datasets are registered to a single calibrator dataset. This process enables users to match corresponding lipid species within and between studies. Formulas (indicated by brackets) are defined in Section 3.1.2. The alignment verification protocols defined in (A) are packaged in RTStaR_Calibrate_v1. The standardization protocols defined in (B) and the registration protocols are packaged in RTStaR_Register_v1 (Supplemental Data).

#### 3.1.1. Defining the calibrator dataset

To determine the theoretical RT of each species in a given dataset, the user defines a calibrator dataset for their particular LC method. This reference (calibrator) dataset is used to (a) quantify and correct the degree of system-level variation in every run and (b) identify runs in which component-level variation (outliers) precludes alignment. The calibrator dataset must meet two basic criteria:

1. The dataset [after alignment and outlier removal (Fig. 1A)] must be composed of a minimum of 10 independent MS runs.
2. Analytes must be present in at least 66 % of all spectra after alignment and outlier exclusion for that particular species to be analyzed. RTStaR_Calibrate automatically removes species that do not meet the 66 % criterion from the calibration list.

#### 3.1.2. Aligning the calibrator dataset

The internal lipid references (IRs) added to span the RT region of interest in the calibrator dataset are assessed first (Fig. 1A, left panel). The number of IRs required must be pre-determined by the user. Our laboratory uses a targeted lipidomics discovery approach to establish this number for each new lipidome. We analyze a small cohort (e.g., n=2-5) of new organisms and/or new sample origins in precursor ion scan (PIS) or neutral loss (NLS) mode, profiling the lipidome of interest. The number of peaks (assuming the best-practice quality control criteria defined in Supplementary Data), their m/z, the number of isobaric peaks, and the RT span of the target m/z range are recorded. Using these data, the number of IRs required to span the RT range of interest (approximately one IR per RT min in the LC methodologies tested) is calculated. Lipid abundances are subsequently determined in multiple reaction monitoring (MRM) by including all the m/z transitions necessary to profile the complete lipidome of interest and the required IRs. Once MRM data for an entire calibrator study is collected, the theoretical IR RTs in the dataset are then established by calculating their respective means (Fig. 1A, left panel).

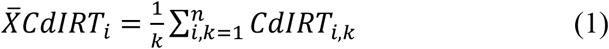

where 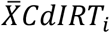 denotes the RT mean of i^th^ IR in all the runs of the calibrator dataset and *CdIRT_i,k_*, the empirical RT of the i^th^ IR in the k^th^ run. *i* includes both the number of IRs added at time of extraction and the number of IRs added at time of MS.

Next, the system-level variation from the theoretical mean for each IR in every run is calculated as the distance (in min) from the 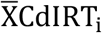.

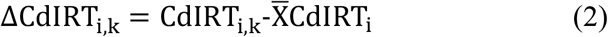

where ΔCdIRT_i,k_ is the distance of the i-th IR species in the k-th run from the mean RT of all runs.

The calibrator equation for each run is defined as:

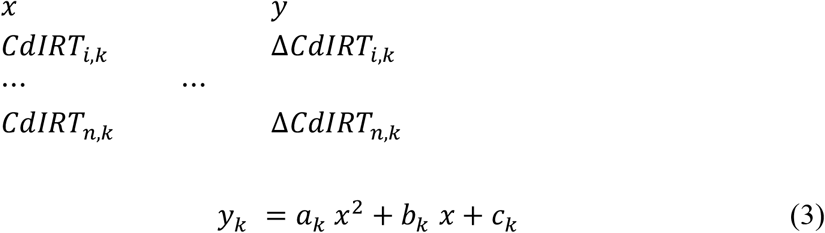

where *a_k_*, *b_k_*, and *c_k_* are the specific parameters of a run calculated by fitting a quadratic regression that relates all of the empirical IR RTs to their respective distances from their theoretical means.

These specific parameters are used to recalculate the aligned RT of each IRs.

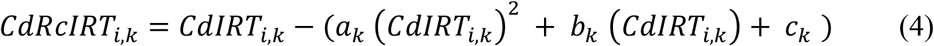

where *CdRcIRT_i,k_* denotes the recalculated (aligned) RT of the i^th^ IR in the k^th^ run. These recalculated IR values, *CdRcIRT_i,k_*, are used to assess the degree of component-level variation in each spectra. RTStaR cannot align runs with extreme component-level variations. To identify these unalignable runs, the recalculated IRs are analyzed for outliers using the Robust regression and Outlier removal (ROUT) method (Motulsky and Brown, 2006). Q values are set to 1 %. The empirical RT values for each outlier species are then re-examined by the operator. If any data acquisition errors (i.e., human error inputting the wrong RT value) are detected, the values are corrected and the IR RTs are iteratively recalculated using Equation (1 to 4). If no human errors are detected, the *CdRcIRT_i,k_* is considered a *bona fide* outlier. In this case, the component-level variation in that particular run is said to exceed the capacity of RTStaR for alignment. The entire run is discarded from the calibrator dataset and the average IR RTs 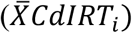 are recalculated. The process is reiterated until no IR RT outliers are detected. Thus, manual and component-level errors are easily flagged by this “go-no-go” outlier statistic and quickly corrected or excluded. The final calibrator dataset must contain a minimum of 10 runs after outlier runs are discarded.

Once the IR RT run-dependent parameters *a_k_*, *b_k_*, and *c_k_* are obtained through Equation (3), the RTs of each species (*CdSRT_m,k_*) are then standardized by adapting Equation (5) (Fig. 1A, right panel).

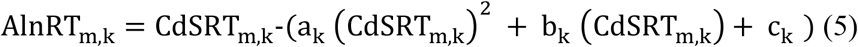

where AlnRT_m,k_ denotes the aligned (theoretical) RT and CdSRT_m,k_, the empirical RT of the m^th^ species in the k^th^ run.

Data acquisition errors are monitored through a non-iterative outlier identification process. At this stage, only outlier *species* (not *runs*) are discarded as the IRs (and thus spectra) have already met criteria. Following a single outlier exclusion assessment using the ROUT method (Motulsky and Brown, 2006), we apply a second statistical criteria, the Median Absolute Deviation (MAD) method (Leys, et al., 2013), to assure that the elution time differences (i.e., component-level variations) between AlnRT_m,k_ fall within three MAD. Species with log_10_-transformed SD over this threshold are excluded. Finally, any aligned species that is not represented in 66 % of all runs is discarded from the entire dataset. This criterion ensures that there are sufficient replicates to calculate the theoretical aligned RT AlnRT_m,k_.

#### 3.1.3. Using the calibrator dataset to identify corresponding lipids across multiple experimental datasets

It is possible to match corresponding and identify unique lipid species in two or more datasets by standardizing their RTs to that of the calibrator dataset (Fig. 1B). Each dataset must share the same nLC method and, ideally, the same IRs although we show in Supplementary Data that omission of some IRs can be tolerated. Critically, *only species that elute within the range of the IRs held in common between datasets can be effectively registered*.

In the standardization algorithm (Fig. 1B, left panel), the experimental IR RTs (EdIRT_i,k_) shared between the calibrator and experimental datasets are recalculated (i.e., standardized to the calibrator dataset). This process accounts for system-level variation in each experimental run. First, the means of all recalculated calibrator IR RTs are determined. These values act as anchor points for the RT standardization of the experimental runs.

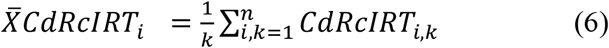

where 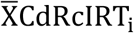 is the mean of the recalculated RT of the i^th^ IR in the k^th^ run of the calibrator dataset.

Next, the system-level variation of every IR RT in the experimental dataset is calculated (Fig. 1B, left panel). Here, RTStaR standardizes one experimental dataset at a time.

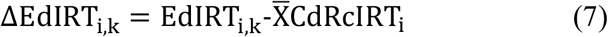

where ΔEdIRT_i,k_ is the distance (in min) of the i^th^ IR species in the k^th^ run from the mean recalculated RT of all runs in the calibrator dataset.

The run-dependent standardized equations are defined as:

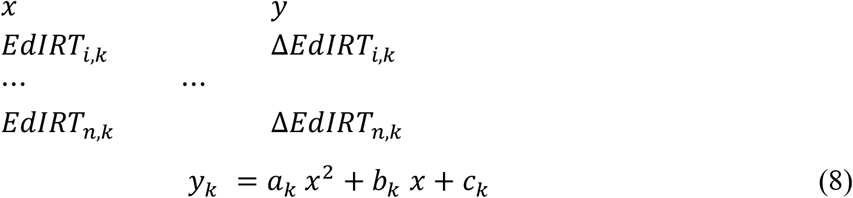

where *a_k_*, *b_k_*, and *c_k_* are the specific parameters of a run calculated by fitting a quadratic regression that relates all of the empirical IR RTs of the experimental dataset to their respective distances from the recalculated mean RTs of the IRs in the calibrator dataset.

These specific parameters (*a_k_*, *b_k_*, and *c_k_*) are used to recalculate the standardized RT of each EdIRT_i,k_.

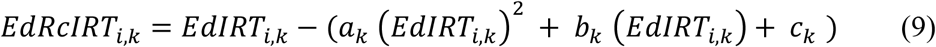

where EdRcIRT_i,k_ denotes the recalculated (standardized) RT of the i^th^ IR in the k^th^ run in the experimental dataset. These recalculated IR values EdRcIRT_i,k_ are used to quality control the nLC of experimental dataset using a process analogous to that described above for calibration. The EdRcIRT_i,k_ values are analyzed for outliers using the ROUT method with a Q set to 1 %. Any EdIRT_i,k_ value identified as an outlier following recalculation as a result of data acquisition errors are corrected and the EdRcIRT_i,k_ are recalculated using Equations 7 to 9. If the RT of an IR in a run is a *bona fide* outlier, the entire run is discarded.

Not all datasets can be used to calibrate different experimental lipidomes. In cases where the system-level variations different between datasets exceed the capacity of RTStaR to register spectra, the algorithm is unable to match corresponding or identify unique species accurately. To verify whether the chosen calibrator can register a given experimental dataset, the user first verifies whether the algorithm accurately matches all of the IRs held in common by both datasets. If experimental IRs are not matched, a different calibrator dataset must be used to align that particular lipidome. This go-no-go criteria is embedded in RTStaR_Register and can be simply established by assessing the Lin’s concordance correlation coefficient between EdRcIRT_i,k_ and its respective 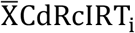 If the lowest Lin’s concordance correlation coefficient in a given run is equal to or greater than 0.9995, all IR RTs will be registered and accuracy for correspondence will be 95% or greater.

Once the registration passes these IR criteria, the experimental species RT (EdSRT_i,k_) are then standardized by adapting Equation 10.

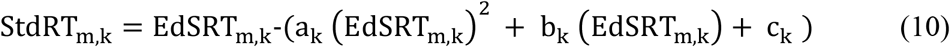

where StdRT_m,k_ denotes the standardized (registered) RT and EdSRT_m,k_, the empirical RT of the m^th^ species in the k^th^ run. Again, data acquisition errors are monitored through outlier identification as described above. Outlier species that cannot be registered in a particular run are flagged and discarded. This process is repeated for each experimental dataset under investigation and corresponding (or novel) lipids present are identified across multiple lipidomes.

## 4 Implementation

### 4.1 Accurate alignment of RTs of 71 unique endogenous metabolites and six IRs in 1001 MS runs

The RTStaR algorithm is packaged as two Excel^™^ workbooks: RTStaR_Calibrate_v1.xlsx and RTStaR_Register_v1.xlsx (Supplementary Data). In addition, workbooks that include two datasets are provided (Supplementary Data). We used a population-based dataset of GPC metabolites extracted from serum of 1001 male and female adolescents representing 84475 analytes with unique or isobaric m/z and 6006 IRs across all runs to establish the calibrator dataset requirements. In this study, chromatographic methodology remained constant but multiple sources of system-level variations were introduced: (1) samples were extracted by eight different investigators; (2) biological replicates were run by three different operators over a period of eight months; (3) MS was run 24 hours a day, 7 days a week until maintenance was required to introduce changes in column performance with repeated usage, etc.; (4) HVAC cooling to the room was turned off and on for a 24 h period twice during the 1001 runs (affecting 24 runs) to model extreme increases and decreases in ambient temperature; (5) packed chromatographic columns were changed six times over the course of the full study; (6) LC capillaries, restriction columns, and electrospray emitters were changed as needed when operators detected component-level variations that necessitated re-running samples (i.e., tubing clogging, random pressure fluctuations etc.); and (7) the nanosource was cleaned once.

Theoretically, lipid species separated by the same chromatography should always elute with identical RTs. Practically, RTs vary dramatically between runs (Fig. 2, Supplementary Data Fig. 1). To address this problem, we applied Equations 1–4 (Fig. 1A), packaged in the first worksheet of RTStaR_Calibrate_v1 (1. Align the Calibrator IRs). Prior to RTStaR alignment, major shifts in empirical RTs of both the endogenous species (gray) and IRs (blue) over the 1001 runs were clearly observed (Fig. 2, left panel). Many of the major variations corresponded to timing of column replacement (Fig. 2, arrowheads). These shifts were effectively corrected following RTStaR alignment (Fig. 2, right panel). Importantly, alignment did not affect the theoretical RT; fold change in the average RT was 0.99-fold (i.e., was equivalent). Only the variance around these mean values were reduced, i.e., the mean standard deviations (SD) decreased by approximately 85-fold across IRs and runs. To quantify the effectiveness of the RTStaR alignment algorithm was quantified, we applied Equation 5 (Fig. 1A), packaged in the second worksheet of RTStaR_Calibrate_v1 (2. Align Calibrator Species). Descriptive statistics of results are presented in the third worksheet of the sample datasets provided (3. Calibrator Results, Supplementary Data).

**Fig. 2.**
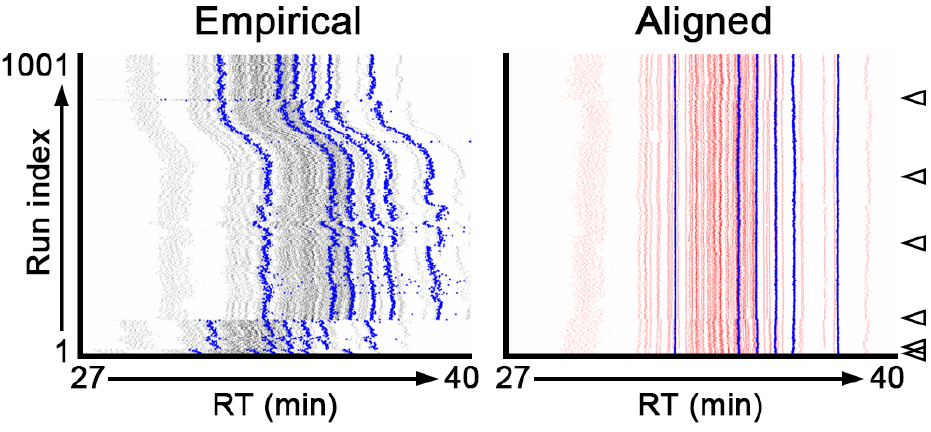
RTStaR alignment of the calibrator dataset. Alignment effectively corrected the system-level variations of endogenous species (grey, empirical; red, aligned) that fell within the range of IR RTs (blue). Empirical run-to-run variations associated not only with column changes (indicated by arrowheads) but also with other unidentified sources were corrected effectively (compare vertical endogenous species alignments between left [gray, empirical] and right [red, aligned] panels) as well as IR alignment (blue).

### 4.2 RTStaR can align isobaric species eluting in close proximity

Minor changes in GPC metabolite structures generate numerous isobaric species that share the same m/z yet elute with proximal (but quantifiably different) RTs (Fig. 3A). In our validation dataset, the 85 endogenous and 6 IR species detected represented 49 distinct m/z. Seventy-three species in each run were isobaric; 40 species were isobaric pairs; 33 species were isobaric triplets. To establish the resolution required to match corresponding species, the distances separating isobaric species were quantified in every run (Fig. 3A, right panel, Supplementary Data). Empirical RTs between isobaric species clearly overlapped from run to run, precluding unambiguous assignation of the majority of species (Fig. 3B, left panel). Only 8 isobaric species, eluting at least 111 sec apart, could be matched (Fig. 3B, left panel). By contrast, all of the isobaric species could be distinguished following RTStaR alignment, i.e., no overlap in SD was observed (Fig. 3B, right panel and inset). To further quantify this marked improvement in discrimination, a resolution equation was applied, modified from (D’Andrea, et al., 2007):

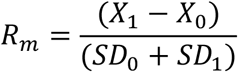

where the numerator is the distance of each downstream isobaric species X_1_ from it upstream X_0_ and the denominator is the sum of the SD for normalized X_0_ and X_1_ RTs (Fig. 3C). RTStaR alignment improved the mean resolution (R_m_) by 33-fold (Fig. 3C). The effect of this registration was demonstrated by overlaying 4 representative XICs pre-(top panel) and post-(bottom panel) RTStaR alignment (Fig. 3D). Unambiguous resolution of a proximal isobaric triplet was evident after algorithmic correction of each cycle time read (Fig 3D).

**Fig. 3.**
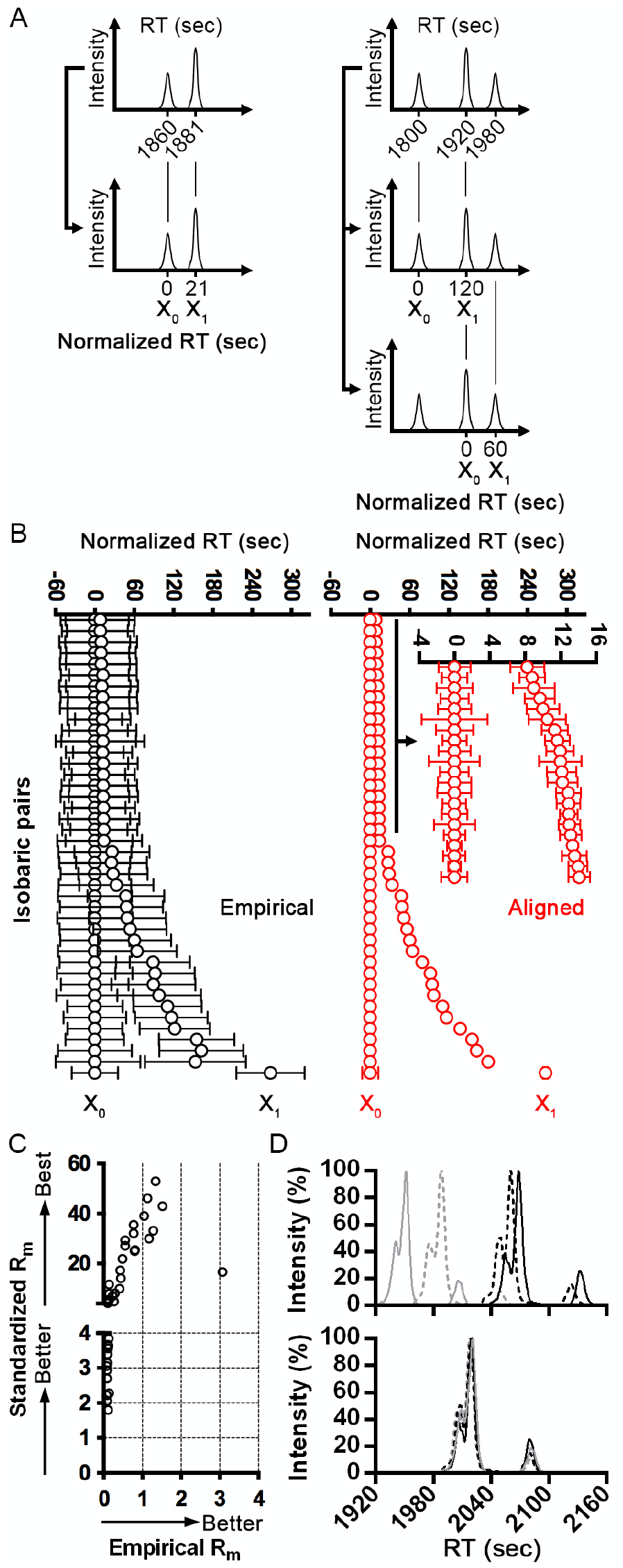
Alignment and identification of corresponding isobaric species in close proximity. (A) Schematic of proximal isobaric pairs (left panel) and triplets (right panel). RT at X_0_ and X_1_ were calculated for each isobaric pair. (B) RTs pre-(left panel) and post-(right panel) RTStaR alignment. Data represent the mean normalized RT for each isobaric X_0_ and X_1_ pair ± SD. Each species was clearly distinguished without overlap of SD after alignment. (C) A higher R_m_ is indicative of a better resolution of isobaric species. R_m_ markedly improved after RTStaR alignment. (D) Extracted-ion chromatogram (XIC) from four samples each containing three isobaric species (482.4 m/z) were aligned using RTStaR. Individual XICs are indicated by black, black-hashed, grey, and grey-hashed lines. The top panel depicts overlay of the empirical XIsC. Bottom panel depicts precise overlay of the XICs following alignment of each cycle time read.

### 4.3 Datasets composed of distinct lipid metabolites can be compared following RTStaR standardization and registration

A useful advance in chromatographic analysis would be to simply yet unambiguously (a) match corresponding lipid species across multiple datasets and (b) identify novel species specific to a given lipidome. RTStaR addresses the problem of lipid correspondence by standardizing the IRs of different experimental datasets to a single calibrator dataset and then registering these standardized datasets to each other. To validate lipidome comparisons, we first anchored the IR RTs of two different profiling studies (mouse NonTg and Tg hippocampal and plasma GPC second messenger lipidomes) to the recalculated calibrator IR RTs 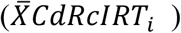 of the human serum calibrator dataset using Equations 6–9 (Fig. 1B). These equations are packaged in the first worksheet of RTStaR_Register_v1 (1. Standardize Experimental IRs). These lipidomes were chosen to challenge RTStaR with diverse experimental effects predicted to influence matrix composition (i.e., sex, treatment, genotype, animal species, tissue, fluids, etc.) All experimental datasets contained the same six IRs. In Supplementary data, we show that RTs of the IRs and endogenous analytes in two different datasets from two different biofluids and species and be reliably standardized to the human serum plasma dataset. An additional useful feature is the capacity to predict the theoretical empirical IR RT (*TheoEmpRT_i,k_*) of all species found in the calibrator dataset in any experimental dataset. This feature allows the user to scan rapidly a new dataset for lipids that correspond with those in the calibrator dataset. This feature is packaged in the seventh worksheet of RTStaR_Register (7. Predict Empirical RTs) and described in Supplementary Data.

### 4.4 RTStaR can match corresponding and identify unique species across lipidomes

To demonstrate that RTStaR could match corresponding and identify new isobaric species across multiple datasets, we first standardized the lipid species present in murine hippocampus and plasma experimental datasets to the human serum calibrator dataset (Fig. 4A,B). We then registered all three datasets (Fig. 4B). The murine hippocampus dataset contained 79 species in each run eluting within the range of the first and last IR RT. Of these 79 species, 66 shared m/z with species in the calibrator dataset. Thirteen species were unique. Of the 66 species potentially held in common with the human serum calibrator dataset, 31 distinct isobaric lipids were represented. In murine plasma, 47 species eluted within the RT range of the IRs. These 47 species shared 22 common m/z with lipids in the calibrator dataset. Isobaric lipids can only be distinguished if the distance between their peaks is measurable. As shown in Fig 3, we were able to resolve all of the isobaric peaks present in the calibrator dataset. However, peak to peak resolution is dependent not only on the quality of chromatography but also the MS cycle time thus capacity to match proximal isobaric species between different datasets is finite (Fig. 4A). Peak to peak distance must exceed machine cycle time specifications by a factor of three for isobaric species to be quantifiably distinguished as separate peaks. For example, the MS cycle time of the MRM methodology we used to analyze the human serum, murine hippocampus, and murine plasma datasets was 0.033 min (Fig. 4A). Thus, the smallest distance between isobaric lipids distinguishable under optimal chromatographic conditions would be 0.099 min or three cycle reads. Practically, peaks must be separated by a distance that exceeds four cycle reads (i.e., > 0.132 min in this particular methodology) considering full-width half-maximum peak values of even the best nLC separation (Fig. 4A). In Figure 3B, we found that that the most proximal isobaric pair in the calibrator dataset, separated by 8.2 sec or 0.137 min, could be accurately aligned and both species statistically separated. Thus, we show empirically that a lipid species present in either the calibrator or experimental dataset can be accurately registered (and thus matched or identified as a unique species) if (a) the difference between their mean standardized 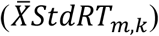 RT in the calibrator dataset and their mean aligned RT 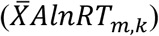 in the experimental dataset is less than twice the cycle time of the particular MS methodology (i.e., ± 0.066 sec in this example) and (b) their m/z is within the MS tolerance (± 1 m/z) (i.e., they share the same m/z). Based on these results, we built these criteria into RTStaR for any user-inputted cycle time or MS tolerance (see RTStaR_Register (5. Identify Corresponding Lipids). The algorithm now automatically excludes isobaric alignments and correspondences when the log_10_ transformed SD is greater than three MAD (see RTStaR_Register (3. Standardize Exp Species and 4. Experimental Results).

**Fig. 4.**
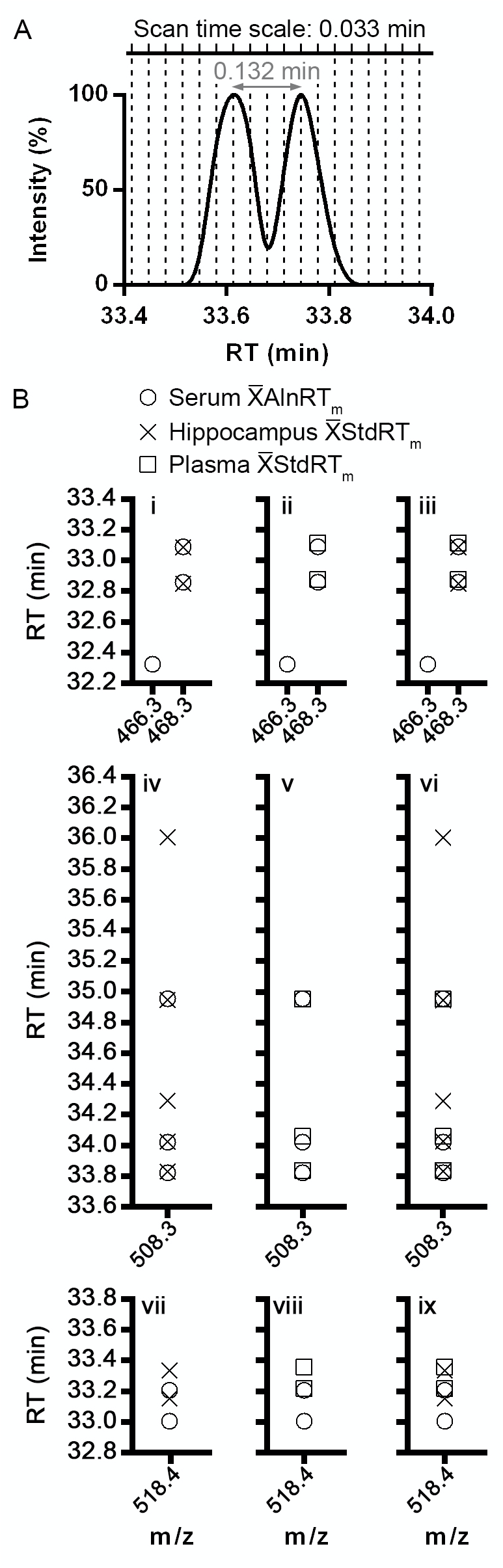
RTStaR can register RTs across multiple datasets, match corresponding isobaric peaks, and identify novel analytes. (A) Schematic of an extracted ion chromatogram depicting two isobaric species and the dependence of resolving these species on MS cycle time. (B) Registration of endogenous species in the human serum, murine hippocampal, and murine plasma datasets after RTStaR alignment and standardization. Data represent 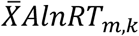 and 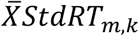 values for lipids with m/z of 466.3, 468.3, 508.3, and 518.4 in human serum (calibrator dataset), murine hippocampal, and murine plasma datasets (experimental datasets).

We show, in the mouse hippocampal dataset, 47 of its 66 GPC metabolites sharing m/z with analytes present in human serum were matches. The remaining 19 species were unique to the hippocampus despite sharing m/z with proximal species in the calibrator dataset. Out of 47 species in the mouse plasma dataset, 37 were matched to the serum calibrator dataset. Ten were specific to murine plasma. Moreover, when we compared the 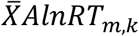 and 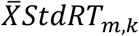 values of all three datasets, we found that we could distinguish between corresponding species common to all three datasets, corresponding lipids found in 2 of 3 datasets, and lipids unique to a single dataset. Figure 4B (panels i-iii) shows the simplest example wherein the human serum dataset contained a unique species at 466.3 m/z not present in either the mouse hippocampal or plasma datasets. All three datasets shared two isobaric species at 468.3 m/z registered with near perfect alignment after RTStaR standardization (Fig. 4B, panel iii). A more complicated example is presented in Fig. 4B (panels iv-vi). The human serum dataset contained three isobaric species with an m/z of 508.3; the murine hippocampal dataset contained five species; the murine plasma dataset contained three species. After standardization, the first, second, and fourth species of the hippocampus dataset matched to the three species found in both the human serum and mouse plasma dataset (Fig. 4B, panels iv-vi). The most complex example is presented Fig 4B (panels vii-ix) wherein novel species eluting in close proximity to corresponding lipids were distinguished. Here, two isobaric species were detected with an m/z of 518.4 in all three datasets. After RTStaR standardization and registration, these species were resolved as three distinct species. The first eluting species was specific to the serum dataset (33.00 ± 0.03 min) (Fig. 4B, panel ix) and while the last was specific to both mouse datasets (hippocampus 33.33 ± 0.04 min and plasma 33.36 ± 0.06 min) (Fig. 4B, panel ix). The middle species was matched between the murine hippocampus (33.15 ± 0.03 min), the murine plasma (33.22 ± 0.05 min), and the human serum dataset (33.21 ± 0.03 min) (Fig. 4B, panels vii-ix). The log transformed SD for these correspondences was less than three MAD and differences between alignments (thus different species) were greater than three MAD. These data demonstrate the capacity of RTStaR to differentiate distinct isobaric doublets between multiple datasets despite their close proximities. RTStaR functions independently of LC and MS methodology as long as datasets being aligned share the same methods.

Lastly, we tested RTStaR’s platform independence with respect to nLC and MS, including legacy MS. We used RTStaR to align human and murine parietal-temporal cortex datasets (a) longitudinally assessing GPC second messenger lipidomes across the lifespan N5 NonTg and TgCRND8 mice, (b) in NonTgPAFR^+/+^, NonTgPAFR^-/-^, TgPAFR^+/+^ and TgPAFR^-/-^ littermates, (c) in post-mortem human brain of Alzheimer’s disease and age-matched human controls. Data were collected using a different nLC protocol on a legacy QTRAP 2000 at a cycle time of 0.028 min with 5 IRs held in common. RTStaR_Register was capable of aligning datasets as long as the *EdRcIRT_i,k_* with its respective 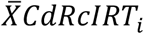 was equal to or greater than 0.9995 (Lin’s concordance correlation coefficient).

## 5 Discussion

Comparative lipidomics lack sufficient bioinformatic capacity to register rapidly and reproducibly corresponding analytes and unambiguously identify unique lipid species unique across lipidomes. To begin to address this bioinformatic need, we developed the RTStaR algorithm. RTStaR combines warp functions, used to estimate the system-level variations particular to a given nLC-nESI-MS/MS methodology in a calibrator dataset, with direct match correspondence functions used to register multiple datasets applying the same LC to this comparator. We show here that (a) the calibrator dataset must be composed of a minimum of 10 independent MS runs; (b) analytes must be present in at least 66 % of all runs after alignment and outlier exclusion; (c) the Lin’s concordance correlation coefficient between *EdRcIRT_i,k_* and its respective 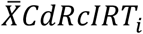 must be equal to or greater than 0.9995 in each MS spectra being registered for a dataset to be used as a calibrator dataset and; (d) sufficient IRs must be introduced in both calibrator and experimental datasets to flank RT range of interest (approximately one IR RT per min). When these criteria are met, RTStaR aligns a given dataset and registers this calibrator dataset with other lipidomes with a correspondence accuracy of 95 % or higher.

## 6 Conclusions

The complete algorithm is packaged in two modular Excel^™^ workbook templates for easy implementation. RTStaR is freely available and can be downloaded from the India Taylor Lipidomics Research Platform website http://www.neurolipidomics.com/resourcesLIPIDS.html.

## Acknowledgements

The following authors are included under the SYS Consortium for patient biosample collection: Michal Abrahamowicz (McGill University), Daniel Gaudet (University of Montreal), Gabriel Leonard (McGill University), Michel Perron (University of Quebec in Chicoutimi), Louis Richer (University of Quebec in Chicoutimi), Jean Seguin (University of Montreal), Suzanne Veillette (CEGEP Jonquiere). We thank Dr Pierre Bel for critical advice. We gratefully acknowledge the beta testing and program revisions provided by Graeme McDowell, Samantha Sherman, Timothea Le, David Bellamy, Cory Lefebvre, and Jun Yu Gao.

