## Supplemental Data for "Retention Time Standardization and Registration (RTStaR): An algorithm that matches corresponding and identifies unique species in nanoliquid chromatography-nanoelectrospray ionization-mass spectrometry lipidomic datasets"

### 1 SUPPLEMENTARY METHODS

#### 1.1 Sample collection and lipid extraction

Human blood was collected in silicone-coated gold BD Hemogard™ tubes (#367986, BD, ON, Canada) from 1001 male and female adolescents, 12 to 18 years of age, enrolled in the Saguenay Youth Study (SYS), a cross-sectional of study of cardio-metabolic and brain health (Paus, et al., 2015). Consent was obtained in strict accordance with the research ethics committees of the Chicoutimi Hospital (Chicoutimi, QC, Canada) and the Hospital for Sick Children (Toronto, ON, Canada). After clotting, serum was separated by centrifugation at 1300 x g for 15 min at room temperature and stored in 1 ml aliquots -80 °C. Parietal-temporal cortex of n=5 patients suffering from Alzheimer's disease and n=5 age-matched controls were obtained from the Douglas Hospital Research Centre Brain Bank (Montreal, QC, Canada) and lipids were extracted as described (Ryan, et al., 2009).

Hippocampi were dissected from 30 six-month-old female N5 NonTg and TgCRND8 N5 C57BL/6 x C3H/He mice (Granger, et al., 2016; Wang, et al., 2013). Blood from these animals was collected in heparin-coated tubes (#022379216, Eppendorf, ON, Canada) and plasma was prepared by collecting the supernatant following centrifugation at 1500 x g for 15 min at 4 °C. Experimental animals received daily dietary supplementation of 75 mg/kg body weight/day omega-3 free fatty acids (40/20 EE 1000 mg fish oil capsules, Ocean Nutrition, NS, Canada) or 75 mg/kg body weight/day omega-6 and omega-9 free fatty acids (1000 mg corn/soybean oil capsules, Ocean Nutrition, NS, Canada) emulsified in vehicle (25 % sweetened condensed milk) for four months. Mice receiving vehicle daily served as controls. In addition, parietal-temporal cortices were dissected from 50 female N5 NonTg and TgCRND8 littermates at two, four, six, eight, twelve or eighteen months of age and 49 male and female N5 NonTgPAFR$^{+/+}$, NonTgPAFR$^{-/-}$, TgCRND8/PAFR$^{+/+}$, and TgCRND8/PAFR$^{-/-}$ littermates between two and eight months of age. Dissections were performed as previously described (Wang, et al., 2013). Tissue was flash frozen in liquid nitrogen and stored at -80 °C until extraction. Experiments were approved by the Animal Care Committee of the University of Ottawa and performed in strict accordance with the ethical guidelines for experimentation of the Canadian Council for Animal Care.

#### 1.2 Internal reference (IR) species and lipid standards dataset

Up to six IRs, PC(13:0/0:0) #855476, PC(12:0/13:0) #LM-1000, $D_4$-PC(O-16:0/0:0) #360906, $D_4$-PC(O-16:0/2:0) #360900, $D_4$-PC(O-18:0/0:0) #10010228, and $D_4$-PC(O-18:0/2:0) #10010229, spanning a mass-to-charge ratio (m/z) range of 454.3 to 636.5, were added to each sample. PC(13:0/0:0) and PC(12:0/13:0) were from Avanti Polar Lipid (AL, USA). Deuterated species and PC(O-16:0/O-2:0) (#78858-42-1) were from Cayman Chemical (MI, USA). The non-naturally occurring PC(13:0/0:0) and PC(12:0/13:0) were added at 187.3 ng and 500 ng per sample respectively at time of extraction. Deuterated isotopologues of four naturally occurring PC metabolites were added at time of nLC-nESI-MS/MS as a mixture containing 1.25 ng of $D_4$-PC(O-16:0/2:0), 1.25 ng of $D_4$-PC(O-18:0/2:0), 2.5 ng of $D_4$-PC(O-16:0/0:0) and 2.5 ng of $D_4$-PC(O-18:0/0:0). In the standards dataset, 1.4 µM PC(13:0/0:0), 2.0 µM PC(14:0/0:0) (#110684), 2.0 µM PC(15:0/0:0) (#855576), 2.0 µM PC(16:0/0:0) (#110685), 2.0 µM PC(17:0/0:0) (#110686), 2.0 µM PC(18:0/0:0) (#110687), 2.0 µM PC(22:0/0:0) (#855779), 2.0 µM PC(17:1/0:0) (#110905), 2.0 µM PC(18:1/0:0) (#110688), 2.0 µM PC(P-16:0/0:0) (#110693), 2.0 µM PC(P-18:0/0:0) (#110694), 3.1 µM PC(O-16:0/2:0) (#110853), 3.1 µM PC(O-17:0/2:0) (#110626), 3.1 µM PC(O-18:0/2:0) (#110627), 2.0 µM PC(O-16:0/4:0) (#878115), 2.0 µM PC(O-18:0/4:0) (#878116), 2.0 µM PC(O-16:0/O-2:0) and 1 µM PC(12:0/13:0) were analyzed. Although m/z does not need to be considered when choosing the IRs for RT alignment, best practice routinely includes IR species from different lipid subfamilies (i.e., PC with hydrocarbon chains of different lengths, linkages, or degrees of unsaturation). This enables the same IRs to be used after RTStaR alignment for subsequent normalization and species abundance calculations.

#### 1.3 nLC-nESI-MS/MS

In the development and validation datasets, lipids were prepared for nLC-nESI-MS/MS as follows: 5 µl of lipid extract (or 5 ul of lipid standards) and 2.5 µl of the isotopologue IR master mix were added to 15.75 µl of $H_2O$ (#9831-03, JT Baker, NJ, USA) containing 0.1 % (v/v) formic acid (#14265, Sigma-Aldrich, ON, Canada) and 10 mM ammonium acetate (#2145, OmniPur, ID, USA). Gradient chromatographic separation was performed on a 10 cm × 75 µm (internal diameter) analytical column packed with ReproSil-Pur 120 $C_4$, 5 µm, 120 Å beads (#r15.4e, Dr. Maisch GmbH, Germany) using an Agilent 1100 LC System (Agilent Technologies, CA, USA) (aka nanobore LC running to mimic nLC conditions). The injection volume was 3 µl. The mobile phase consisted of solvent A ($H_2O$) and B (5:2, acetonitrile:isopropanol, v/v) (#A416-4, Fisher Scientific, ON, Canada, #9829-03C, JT Baker, respectively) both containing 0.1 % (v/v) formic acid and 10 mM ammonium acetate (#2145, OmniPur, ID, USA). The sample was loaded for 15 min in 5 % solvent B. Gradient elution was performed with a linear increase of solvent B from 5 to 35 % in one minute and from 35 to 100 % in 14 min. One hundred percent solvent B was maintained for an additional 14 min before the column was equilibrated back to 5 % B for 15 min. The eluent was nanoelectrosprayed into a QTRAP 5500 (SCIEX, MA, USA) through an emitter column (#FS360-50-15-N-20, PicoFrit, New Objective, MA, USA) and analyzed in multiple reaction monitoring (MRM) mode. The parent ion m/z were set in Q1, keeping the Q3 mass analyzer set to 184 m/z detecting the product ion that defines PC. MS parameters were as follows: ion spray voltage (IS), 3500 V, ion source gas 1 (GS1), 10 psi; turbo gas (GS2), 0 psi, curtain gas (CUR), 20 psi; collision gas (CAD), 10 psi, declustering potential (DP), 100 V, entrance potential (EP), 10 V, collision cell exit potential (CXP), 9 V, collision energy (CE), 47 eV, and scan time, 1.98 sec/cycle. High purity nitrogen (#UN1977, Boc Gases, ON, Canada) was used for CUR, GS1, and CAD. Data were acquired using SCIEX Analyst v1.5.1 and analyzed using MultiQuant v2.0.2.

Three additional datasets were acquired on a legacy QTRAP 2000 using a different nLC (nanobore) method. Gradient chromatographic separation was performed using an Agilent 1100 LC System on a 5 cm × 200 µm (ID) analytical column packed with Magic $C_{18}$ AQ 100A 5U 5 µm resin (#PM5/61100/00, Michrom BioResources, CA, USA) coupled to a 5 cm packed analytical emitter column (#PF360-75-10-N-5, PicoFrit, New Objective) packed with the same beads. The injection volume was 8 μl. The mobile phase consisted of $H_2O$ containing 0.1 % formic acid (solvent A) and acetonitrile (#A416-4, Fisher Scientific, ON, Canada) containing 0.1 % formic acid (solvent B). Gradient elution was performed with a linear increase of solvent B from 5 to 30 % in 2 min, 30 to 60 % in 7 min, 60 to 80 % in 33 min, and 80 to 95 % in 4 min. The analytes were nanoelectrosprayed at 3500 volts into a QTRAP 2000 and analyzed in PIS mode for the detection of PCs. CE was set to 40 eV. Data acquisition (m/z and RT) was performed and analyzed using Analyst v1.4.2.

#### 1.4 nLC-nESI-MS/MS quality control criteria

As with any bioinformatic application, spectra must meet best-practice nLC-nESI-MS/MS quality control criteria prior to submission to RTStaR. First, accurate peak identification requires an adequate signal-to-noise ratio of, at least, ten fold (Kim, et al., 2007). Second, peak shape must be relatively symmetrical and smooth to establish reproducibly the point of maximum intensity defining the RT (Barwick, et al., 2006). Third, a sufficient number of IRs must be added either at time of sample extraction, or time of nLC-nESI-MS/MS injection, or both. This number is defined empirically by the span (in minutes) of the RTs of all species of interest present in a given lipidome. We show below (4. Supplementary Results) that IR RTs should be relatively equally distributed across the range of target RTs in both calibrator and experimental datasets and that optimal alignment is achieved when runs contain (a) approximately one IR per minute of RT and (b) the first and last IR RT flank the RTs of all of the endogenous species of interest.

#### 1.5 Calculation of the resolution required to match corresponding isobaric species

To calculate the resolution required to match isobaric species, the distances separating isobaric pairs were quantified in every run. Distances were defined as the peak-to-peak (RT) elution time in sec. For each species, we normalized both the empirical RTs ($CdSRT_{m,k}$) and the aligned RTs ($AlnRT_{m,k}$) setting the isobaric species eluting first to 0. When more than two isobaric species was detected at a given m/z, once the first distance was established, $X_0$ was set to the RT of the second species and $X_1$ was set to the RT of the third (see Fig. 3A, right panel, main text).

Empirical RT:

$$X_0 = CdSRT_{X_0,k} - \bar{X}CdSRT_{X_0}$$
$$X_0 = CdSRT_{X_0,k} - \bar{X}CdSRT_{X_0}$$

Align RT:

$$X_0 = AlnRT_{X_0,k} - \bar{X}AlnRT_{X_0}$$
$$X_1 = AlnRT_{X_1,k} - \bar{X}AlnRT_{X_0}$$

where $X_0$ denotes the isobaric species eluting first and $X_1$ its closet subsequent neighbour (see Fig. 3A, right panel, main tet).

### 2 SUPPLEMENTARY RESULTS

#### 2.1 The RTs of 71 unique endogenous metabolites and six IRs in 1001 MS run can be accurately aligned using RTStaR

Prior to RTStaR alignment, we found that maximal variations in the six IR RTs of the human serum dataset (1001 spectra) ranged from 6.16 min for PC(13:0/0:0) to 9.42 min for PC(12:0/13:0) (Supplementary Fig. 1). After alignment, the maximal RT variation ranged from 0.03 min for PC(13:0/0:0) to 0.13 min for $D_4$-PC(O-18:0/2:0) (Supplementary Fig. 1). Alignment reduced the average minimum to maximum variation by approximately 116-fold across all IRs. Next, we used cumulative frequency graphs to assess the distribution of the IR RTs across runs (Supplementary Fig. 1). Prior to alignment, bi- and tri-modal distributions were

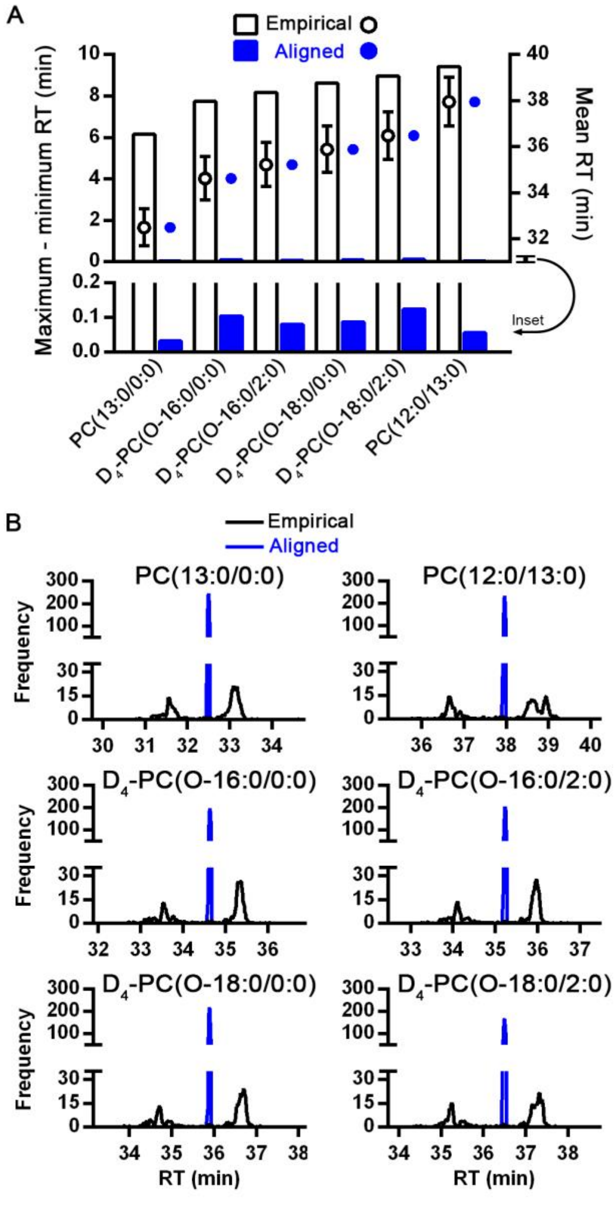


**Supplementary Fig. 1. RTStaR alignment of the calibrator dataset.** (A) Alignment reduced maximum to minimum run-to-run variation in IR RTs (compare white and blue bars, left Y-axis) and their standard deviations around the theoretical IR RT (compare white and blue symbols representing average IR RT ± SD, right Y-axis). In the lower panel (inset), the left Y-axis is expanded between 0 and 0.2 min to display the markedly reduced maximum to minimum run-to-

run variation after calibration. (B) Before alignment, bi-modal, and sometimes tri-modal, RT distributions were detected; indicative of different system-level variations in groups of runs (black lines). After alignment, these differences were effectively resolved, transforming five of six IR RT datasets into Gaussian distributions (blue lines). The exception was the $D_4$-PC(O-16:0/2:0) RT distribution, transformed from a multi-modal to single distribution but with a statistically significant skew ($p < 0.0001$). Statistics were D'Agostino-Pearson omnibus normality tests with $p$ set to 0.05.
observed, likely indicative of different system-level variations in groups of runs. RTStaR calibration transformed the majority of the multi-modal distributions (5 of 6 IRs) into Gaussian distributions (Supplementary Fig. 1).

### 2.2 To compare lipidomes, the RTs of the IRs and endogenous analytes in two different datasets are standardized to the calibrator dataset

To compare lipidomes, each experimental dataset is first standardized separately to the calibrator dataset. Here, we compared mouse hippocampal and plasma lipidomes to the human serum calibrator dataset. The mean of each recalculated calibrator IR RT ($\bar{X}CdRcIRT_i$ ) was determined (Equation 6, Fig. 1B). The relationship between these calibrator anchor points and the IR RTs in experimental run was then established. These parameters were used to standardize the IR RTs across datasets ($EdIRT_{i,k}$), verify the quality of the spectra, and exclude runs that could not be standardized due to extreme component-level variation (Equation 7-9, Fig. 1B). In both experimental datasets, no outliers were detected in the IR RTs; no run was excluded. Supplementary Figure 2A presents the results of the IR RT standardization. Note the near perfect registration of the experimental IRs in both datasets following RTStaR standardization (Supplementary Fig. 1A). Deviation between empirical IR RT ($\bar{X}EdIRT_i$ ) and the $\bar{X}CdRcIRT_i$ was reduced by 96.6 % in mouse hippocampal and 99.5 % in mouse plasma following standardization ($EdRcIRT_{i,k}$) (Supplementary Fig. 1A). Moreover, RTStaR standardization transformed all of the IR RT frequency distributions in both experimental datasets into Gaussian distributions (D'Agostino-Pearson omnibus normality tests with p set to 0.05). Compared to the calibrator dataset, the mouse hippocampal and plasma datasets displayed Lin's concordance correlation coefficients that ranged between 0.9995-0.9999 and 0.9998 to 0.9999 across all runs respectively. Statistics are presented in sample workbook RTStaR_Register_SerumCd_vs_ PlasmaEd.xlsx, '1. Standardize Experimental IRs', "Descriptive Statistics" (Supplementary Data).

The experimental species RTs were then standardized ( $StdRT_{m,k}$ ) using Equation 10 packaged in the second worksheet of RTStaR_Register (2. Standardize Exp Species). Results are found in RTStaR_Register (3. Calibrator Results). In the mouse hippocampal dataset, 35 of 2863 empirical species RT ($EdSRT_{m,k}$) were identified as *bona fide* outliers (data not shown). In the mouse plasma dataset, 35 of 2248 empirical species RT ($EdSRT_{m,k}$) were identified and removed as outliers in RTStaR_Register (3. Calibrator Results). Following outlier identification and standardization, a marked reduction in variance was observed in 85.6 % and 74.4 % of the PC metabolites detected in the murine hippocampus and plasma, respectively (Supplementary Fig. 1B, black symbols). Species alignment, however, *deteriorated* in 14.4 % and 25.6 % of the hippocampal and plasma lipids respectively, meaning that RTStaR standardization *increased* SD around the respective means of these particular lipids. In each case, affected species eluted with a $StdRT_{m,k}$ that fell outside of the range of the recalculated calibrator IR RTs ($EdRcIRT_{i,k}$, Supplementary Fig. 1B, outside). To explore this deviation in more detail, we compared the mean distance that each PC metabolite eluted from its closet IR anchor point to the SD of their $StdRT_{m,k}$ (Supplementary Fig.

1C). Clearly, when the $StdRT_{m,k}$ fell within the range of IR RT span, RTStaR alignment reduced the variance across runs (Supplementary Fig. 1B,C, inside), however if the $StdRT_{m,k}$ was not flanked by an IR anchor point then RTStaR standardization failed (Supplementary Fig. 1B,C, outside). Species that eluted after the last IR were the most affected (Supplementary Fig 1B,C, after). Collectively, these data show that RTStaR can effectively register multiple datasets only if the RTs of target species fall within the RT range of the IR anchor points. Lipids that elute outside of this range are not accurately calibrated.

#### 2.3 The RTs of the IRs and endogenous analytes in two different datasets can be standardized to the calibrator dataset

Each experimental dataset was standardized separately to the calibrator dataset. The mean of each recalculated calibrator IR RT ($\bar{X}CdRcIRT_i$ ) was determined (Equation 6, Fig. 1B). The relationship between these calibrator anchor points and the IR RTs in experimental run was then established. These parameters were used to standardize the IR RTs across datasets ($EdIRT_{i,k}$), verify the quality of the spectra, and exclude runs that could not be standardized due to extreme component-level variation (Equation 7-9, Fig. 1B). In both experimental datasets, no outliers were detected in the IR RTs; no run was excluded. Supplementary Figure 2A presents the results of the IR RT standardization. Note the near perfect registration of the experimental IRs in both datasets following RTStaR standardization (Supplementary Fig. 2A). Deviation between empirical IR RT ($\bar{X}EdIRT_i$ ) and the $\bar{X}CdRcIRT_i$ was reduced by 96.6 % in mouse hippocampal and 99.5 % in mouse plasma following standardization ( $EdRcIRT_{i,k}$ ) (Supplementary Fig. 2A). Moreover, RTStaR standardization transformed all of the IR RT frequency distributions in both experimental datasets into Gaussian distributions (D'Agostino-Pearson omnibus normality tests with p set to 0.05). Compared to the calibrator dataset, the mouse hippocampal and plasma datasets displayed Lin's concordance correlation coefficients that ranged between 0.9995-0.9999 and 0.9998 to 0.9999 across all runs respectively. Statistics are presented in sample workbook RTStaR_Register_SerumCd_vs_ PlasmaEd.xlsx, '1. Standardize Experimental IRs', "Descriptive Statistics" (Supplementary Data).

The experimental species RTs were then standardized ( $StdRT_{m,k}$ ) using Equation 10 packaged in the second worksheet of RTStaR_Register (2. Standardize Exp Species). Results are found in RTStaR_Register (3. Calibrator Results). In the mouse hippocampal dataset, 35 of 2863 empirical species RT ($EdSRT_{m,k}$) were identified as *bona fide* outliers (data not shown). In the mouse plasma dataset, 35 of 2248 empirical species RT ($EdSRT_{m,k}$) were identified and removed as outliers in RTStaR_Register (3. Calibrator Results). Following outlier identification and standardization, a marked reduction in variance was observed in 85.6 % and 74.4 % of the PC metabolites detected in the murine hippocampus and plasma, respectively (Supplementary Fig. 1B, black symbols). Species alignment, however, *deteriorated* in 14.4 % and 25.6 % of the hippocampal and plasma lipids respectively, meaning that RTStaR standardization *increased* SD around the respective means of these particular lipids. In each case, affected species eluted with a $StdRT_{m,k}$ that fell outside of the range of the recalculated calibrator IR RTs ($EdRcIRT_{i,k}$, Supplementary Fig. 2B, outside). To explore this deviation in more detail, we compared the mean distance that each PC metabolite eluted from its closet IR anchor point to the SD of their $StdRT_{m,k}$ (Supplementary Fig. 2C). Clearly, when the $StdRT_{m,k}$ fell within the range of IR RT span, RTStaR alignment reduced the variance across runs (Supplementary Fig. 2B,C, inside), however if the $StdRT_{m,k}$ was not flanked by an IR anchor point then RTStaR standardization failed (Supplementary Fig. 2B,C, outside). Species that eluted after the last IR were the most affected (Supplementary Fig 2B,C, after).

Collectively, these data show that RTStaR can effectively register multiple datasets only if the RTs of target species fall within the RT range of the IR anchor points. Lipids that elute outside of this range are not accurately calibrated.

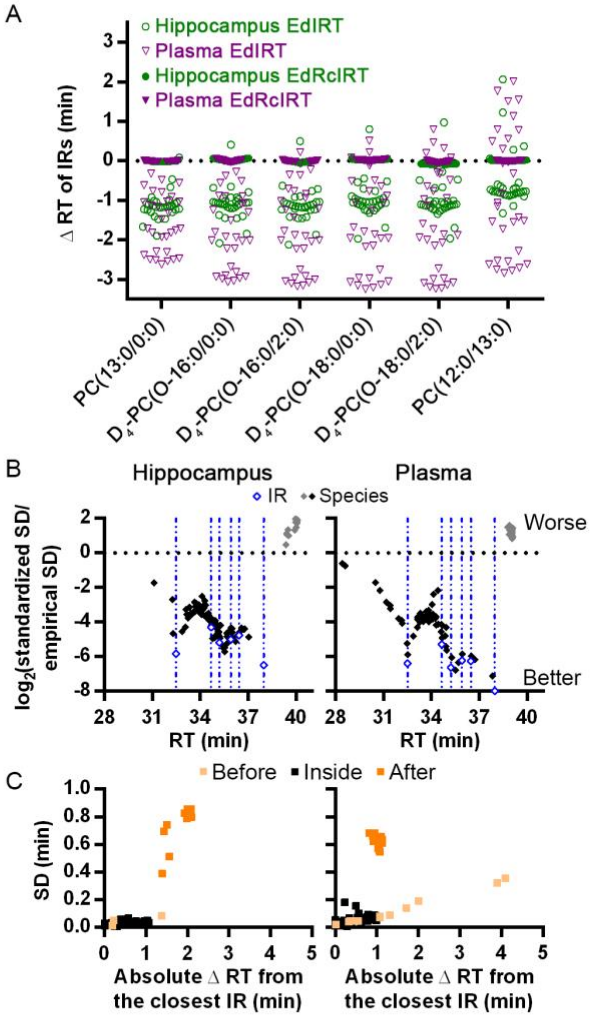


**Supplementary Fig. 2. RTs from multiple datasets can be standardized to the calibrator dataset.** (A) The IR RTs in two distinct lipidomes, a murine plasma and a murine hippocampus PC second messenger datasets, were standardized to the $\bar{X}CdRcIRT_i$ of the calibrator dataset. Open symbols represent the difference in min between the empirical $EdIRT_{i,k}$ of each experimental run and the appropriate $\bar{X}CdRcIRT_i$ of the calibrator dataset. Closed symbols represent this difference after standardization. Note the near perfect registration of the $EdRcIRT_{i,k}$ in both experimental datasets with the calibrator dataset following RTStaR standardization. (B) RTStaR standardization improved the registration of $StdRT_{m,k}$ (black symbols) that eluted within the range of the $EdRcIRT_{i,k}$ (blue symbols, blue lines). RTStaR was less effective at correcting system-level variance that eluted before the initial IR and could not correct the system-level variance for species that eluted after the terminal IR. In these species, SD around the theoretical mean increased following RTStaR standardization (grey symbols). (C) The reduction in variance was clearly dependent upon the distance to the closest IR anchor point (black symbols, inside). Species eluting before the first IR $\bar{X}EdRcIRT_i$ (tangerine symbols, before) were less affected than species than eluted after the last IR $\bar{X}EdRcIRT_i$ (orange symbols, after).

### 2.4 Reverse calculating (i.e., predicting) the empirical RTs facilitates analysis of target analytes and correct peak picking.

A useful feature of the RTStaR algorithm is the capacity to predict the theoretical empirical IR RT ($TheoEmpRT_{i,k}$) of all species found in the calibrator dataset in any experimental dataset. Using RTStaR_Register (7. Predict Empirical RTs), the users are provided with the predicted empirical RT by querying the run specific value reported in RTStaR_Register (7. Predict Empirical RTs).

To obtain the predicted $EdIRT_{i,k}$ and $EdSRT_{m,k}$ values, RTStaR uses the aligned RTs of the calibrator dataset ($AlnRT_{m,k}$):

$$\bar{X}AlnRT_m = \frac{1}{k}\sum_{i,k=1}^{n} AlnRT_{m,k} \quad (11)$$

where $\bar{X}AlnRT_m$ is the mean of the aligned RT of the m$^{th}$ species in the calibrator dataset. As every run must be standardized individually, only the IR empirical RTs of each runs of interest need to be acquired and processed through Equations 7-9 as described in Section 3.2.3 to obtain the run parameters $a_k$, $b_k$, and $c_k$. $\bar{X}AlnRT_m$ is then used to calculate the theoretical RT of that species in a given run.

$$\frac{(b_k - 1) + \sqrt{(b_k - 1)^2 - 4(a_k)(c_k + \bar{X}AlnRT_m)}}{2(-a_k)} = TheoEmpRT_{m,k} \quad (12)$$

where $a_k$, $b_k$, and $c_k$ are obtained from Equations 7-9 and $\mathrm{Theo}EmpRT_{m,k}$ is the theoretical empirical RT of the m$^{th}$ species in the k$^{th}$ run. We show below that the accuracy of this prediction requires that the calibrator and experimental dataset IRs display a Lin’s concordance correlation coefficient of 0.9995 or greater.

We assessed the effectiveness of this standardization in percent (E%) by adapting the equations developed by (Lantos, et al., 2000):

$$E\% = 100 - \left[ABS\left(\frac{EdIRT_{m,k} - TheoRT_{m,k}}{EdIRT_{m,k}}\right)\right]100 \quad (13)$$

For the analytes in hippocampal dataset standardized to the human serum calibrator dataset, the average E% per species after outlier removal ranged between 99.6 % and 100.0 %. The mean percent effectiveness for the 47 species was 99.9 %.

### 2.5 Defining the requirements of a calibrator dataset: Not all datasets can be calibrator datasets

The calibrator dataset used to validate RTStaR was composed of 1001 runs. To establish the minimum runs required, we repeated the calibration using 30, 15, or 10 consecutive runs randomly chosen from the larger population-based dataset. These minimal calibrator datasets were capable of standardizing the mouse plasma dataset with the same results as the population-level calibration when the reproducibility of the alignment of each $EdRcIRT_{i,k}$ with its respective $\bar{X}CdRcIRT_i$ equalling or exceeding a Lin’s concordance correlation coefficient of 0.9995. Based on these results, we established that a calibrator dataset must be composed of a minimum of 10 independent MS runs, after alignment and outlier removal to be effective.

Next, we asked how many IRs must be held in common between a calibrator dataset and an experimental dataset for registration to occur. Using the human serum dataset and its 6 IRs as the calibrator dataset, we sequentially removed IRs from the murine hippocampal dataset and assessed impact on its registration. With all six IRs present in both datasets, 47 PC metabolites in murine hippocampus could be matched with species in human serum (Supplementary Fig. 3A-C, triangles). As expected based on our empirical data (Supplementary Fig 2), when the initial IR with the smallest RT [PC(13:0/0:0)] was excluded, lipid species in the experimental dataset that eluted before this RT could not be matched (Supplementary Fig. 3A, open circle). We then assessed the loss of two consecutive IRs in the middle of the RT range, $D_4$-PC(O-16:0/2:0) and $D_4$-PC(O-18:0/0:0). All of the 47 species were correctly registered, but the average absolute RT distance deteriorated by 9.2 %, from 0.025 to 0.027 min (Supplementary Fig. 3B, open circles). When three consecutive IRs were removed towards the end of the elution range, discarding $D_4$-PC(O-18:0/0:0), $D_4$-PC(O-18:0/2:0) and PC(12:0/13:0) (Supplementary Fig. 3C, open circles), we were unable to register species that eluted after the last IR shared by both datasets. Moreover, registration of the species that remained within the range of common anchor points range deteriorated by 144.3 % with average absolute distance from their calibrator counterparts increasing from 0.025 to 0.060 min. These data demonstrate RTStaR alignment, standardization, and registration function optimally when the calibrator and the experimental datasets share the same IRs. Where IRs differ, only species that fall within the elution range of the RTs held in common can be registered effectively with resolution dependent on the number of IRs present.

Finally, we quantified the reproducibility and the accuracy of RTStaR alignments using different calibrator datasets. We compared inter-calibrator agreement between a dataset composed of 22 lipid standards both calibrating and being calibrated by the human serum dataset (1001 runs), the murine hippocampal dataset (30 runs), and the murine plasma dataset (30 runs). All of the lipids added to the lipid standard dataset were isobaric with endogenous species present in all three lipidomes. By addition of standards to each matrix, we verified that human serum contained 13 of the 20 standards; murine plasma contained 11 of the 20 standards; murine hippocampus contained 13 of the 20 standards. Each lipidome contained at least one corresponding species that differed from one of the other three biological lipidomes. Accuracy of correspondence with these validated results was 100 % when the lowest degree of agreement between $EdRcIRT_{i,k}$ with its respective $\bar{X}CdRcIRT_i$ was equal to or greater than 0.9995 (Lin's concordance correlation coefficient) (Supplementary Fig 3D). Thus, RTStaR requires that (a) a calibrator dataset is composed of a minimum of 10 runs in which each analyte represented in at least 66 % of each spectra, (b) only species that fall between the RTs of calibrator datasets can be effectively registered, and (c) the minimal $EdRcIRT_{i,k}$ concordance with calibrator $\bar{X}CdRcIRT_i$ must equal or exceed a Lin's concordance correlation coefficient of 0.9995 for correspondence accuracy to be 95 % or greater. Here, the user is cautioned that if their calibrator dataset is composed of equal numbers of control and treatment/disease replicates with treatment/disease anticipated to deplete levels of particular analytes below limits of their particular methodology or introduce novel species into a lipidome, they may wish to consider registering experimental to control data using RTStaR Register as these species will not meet the 66 % criteria necessary to establish baseline calibration.

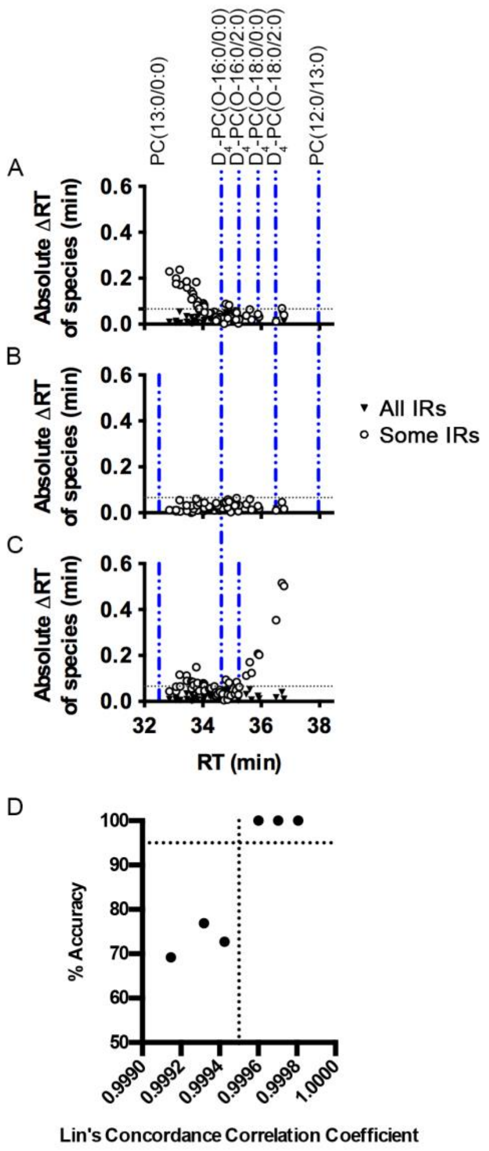


**Supplementary Fig. 3. RTStaR functions optimally when the calibrator and experimental datasets share the same IRs.** (A) Lipid species in the experimental dataset that elute before the first IR held in common between calibrator and experimental datasets cannot be matched to the corresponding lipids present in calibrator dataset. (B) Experimental species that fall within the elution range of the RTs held in common with the calibrator dataset can be registered effectively but with less precision than when all IRs are held in common. (C) Lipids that elute after the terminal IR RT shared by both calibrator and experimental datasets cannot be effectively registered. (D) Accuracy of calibration is dependent on the degree of concordance between the $EdRcIRT_{i,k}$ dataset with its respective $\bar{X}CdRcIRT_i$ . We compared capacity of a dataset composed of 22 standards (Avanti Polar Lipid and Cayman Chemical) to calibrate the human serum, murine plasma, and murine hippocampal datasets and vice versa. The number of endogenous species matching these standards present in each biological lipidomes was verified by addition of exogenous standards. We found that any dataset with a Lin's concordance correlation coefficient of 0.9995 between calibrator $\bar{X}CdRcIRT_i$ and experimental $EdRcIRT_{i,k}$ effectively registered all of the endogenous species correctly with an accuracy of 100 %. Any value below this concordance, failed to reproducibly match corresponding species with an accuracy of greater than 95 %.

### 3 CONTRIBUTIONS

A.P.B. with S.A.L.B. wrote, designed, and validated the RTStaR algorithm. Z.P., T,P., and the Sys Consort collected the human datasets. HX and S.A.L.B co-direct the India Taylor Lipidomic Research Platform in the uOttawa Proteomic Facility where all MS experiments were performed. H.X., Y.W., A.P.B., and S.A.L.B. performed and analyzed the MS experiments with advice from D.F. A.P.B, M.G, and G.P.T. verified platform independence using different nLC methods. S.F. developed and hosts the online resource page as part of the Carleton Immersive Media Studio. A.P.B. with H.X. and S.A.L.B. wrote the manuscript. All authors provided input.
